## Supplementary notes and figures for "Inference of gene regulatory networks using time-series single-cell RNA-seq data with CRISPR perturbations"

### 1 Supplementary notes

#### 1.1 Diagonal elements of A

$\mathbf{A}_{i,i}$ $(i = 1, \ldots, G)$ represents the concentration-dependent effects from gene $i$ to gene $i$ including self-regulation and degradation, which cannot be estimated separately. Elements of $\mathbf{A}^k$ $(k \geq 2)$ include the products of $\mathbf{A}_{i,i}$s. For example, $\mathbf{A}_{i,j} \cdot \mathbf{A}_{i,i}$ $(i \neq j)$ represents the strength of the regulation from gene $j$ to gene $i$ multiplied by the strength of degradation or self-regulation of gene $i$. As only genes $i$ and $j$ are involved, this term should be interpreted as a first-order regulation and not a second-order regulation. Therefore, a more precise explanation for the elements of $\mathbf{A}^k$ $(k \geq 2)$ is that they are a mixture of the effects of $k'$-th order regulation $(k' \leq k)$ and degradation.

#### 1.2 Possible future extensions of RENGE

In this study, RENGE was applied to scCRISPR data for hiPSC, a cell type in a steady state. However, GRN inference can also be performed using scCRISPR data for cells in a transient state of dynamics such as differentiating cells under the assumption that the gene regulation $\mathbf{A}$ remains the same during the dynamics. In this situation, with the gene expression data of wild-type and knockout cells sampled at multiple time points, the GRN can be inferred using the same model equation as in this study. Thus, it is possible, for example, to infer the GRN in cells during differentiation.

The inference accuracy of GRNs using our data could be improved as follows: (1) Use of pesudotime. RENGE is a time series model that uses the sampling time of cells as temporal information. Alternatively, we may develop a model that integrates the information of pseudotime, which is estimated from the expression change in each cell [1], as temporal information with a higher time resolution. (2) Use of data from other modalities. The RENGE method can only infer GRNs from transcriptome data. However, GRN inference methods that utilize the DNA binding motifs of TFs [2, 3], or chromatin accessibility data [4, 5, 3], as well as transcriptome data, have recently been developed and have been proven useful. Therefore, GRN inference methods that integrate time-series transcriptome data following gene perturbations with data from other modalities could improve inference accuracy.

### 2 Supplementary figures

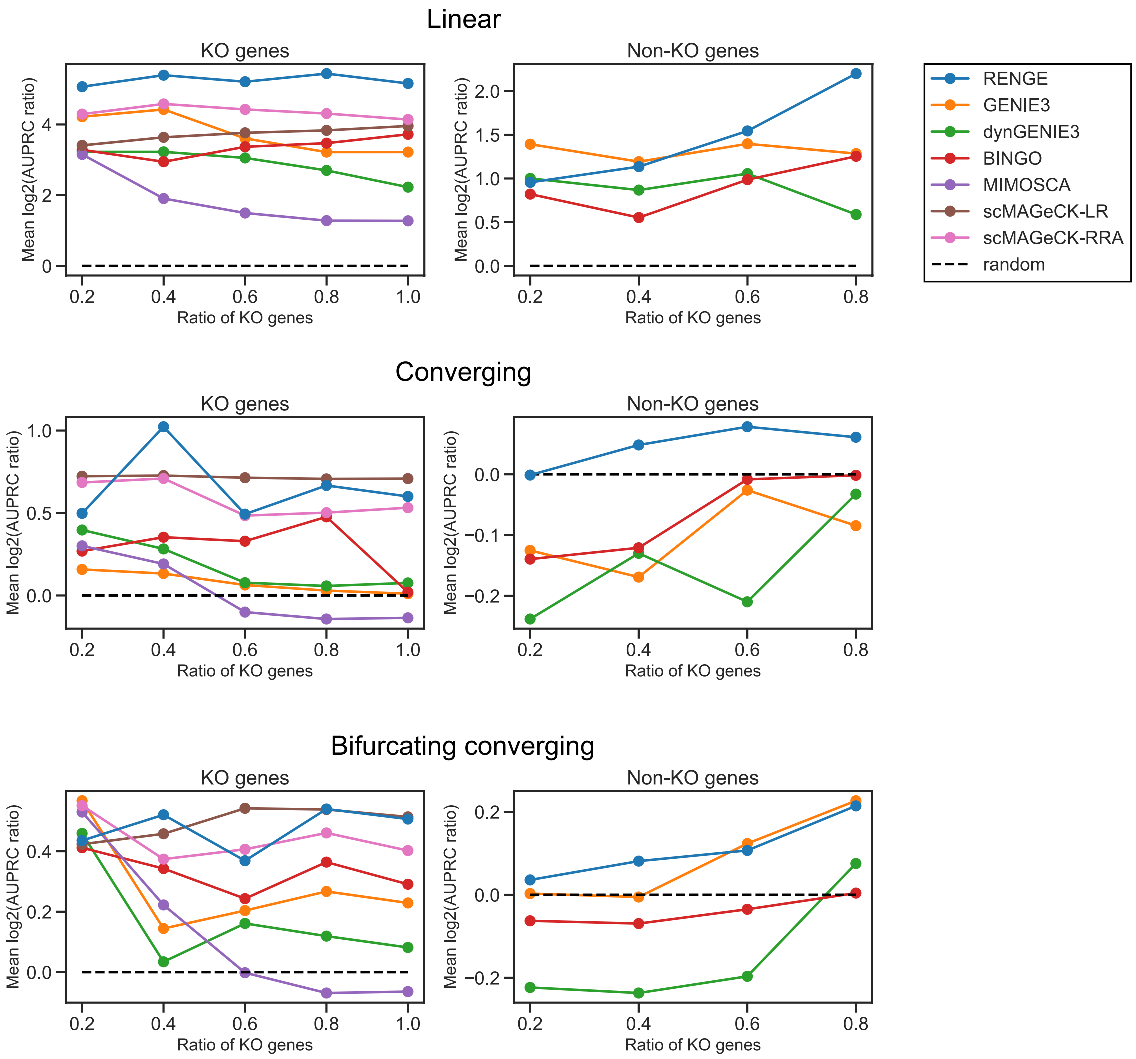


Supplementary Figure 1: **Benchmark results from the simulated data sets for each network generated using the three backbones defined in dyngen.** Horizontal axis: knocked-out gene ratio in the GRNs.

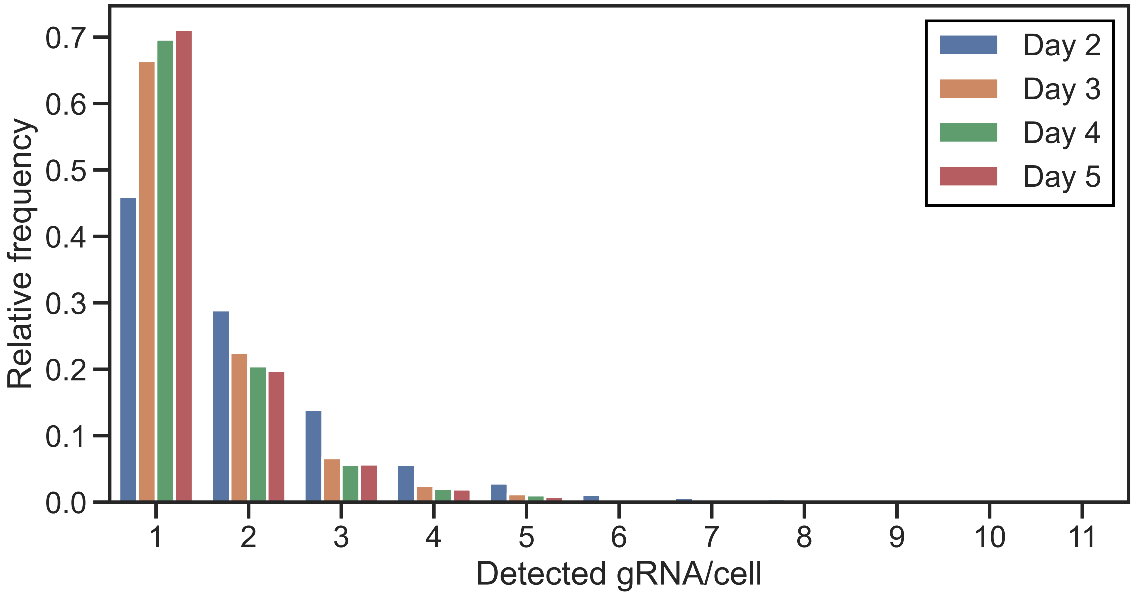


Supplementary Figure 2: **Distribution showing the number of gRNAs detected each day per hiPSC.**

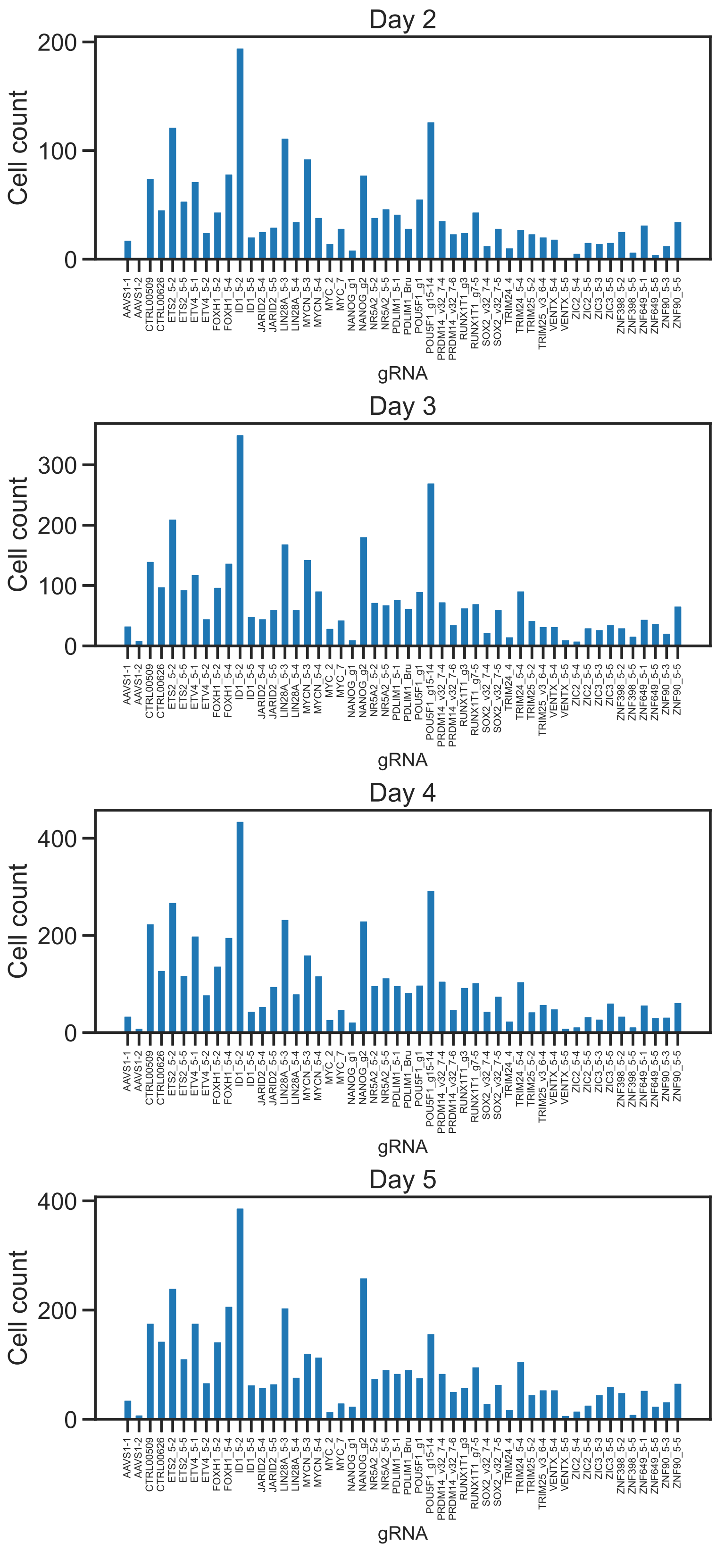


Supplementary Figure 3: **Number of cells in which each gRNA was detected.** Only cells bearing a single gRNA were counted.

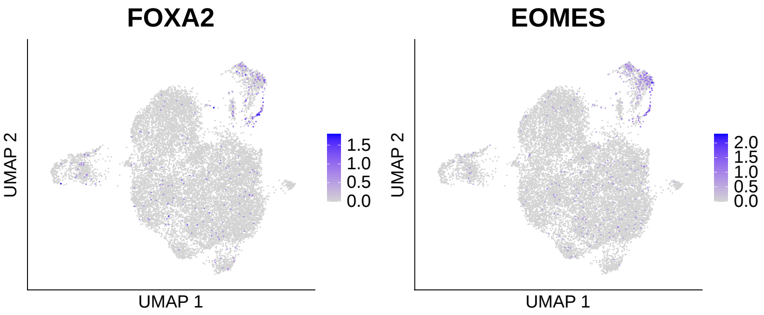


Supplementary Figure 4: **Marker expression on the UMAP plot.** Colors indicate the expression of FOXA2 or EOMES (mesoendoderm markers).

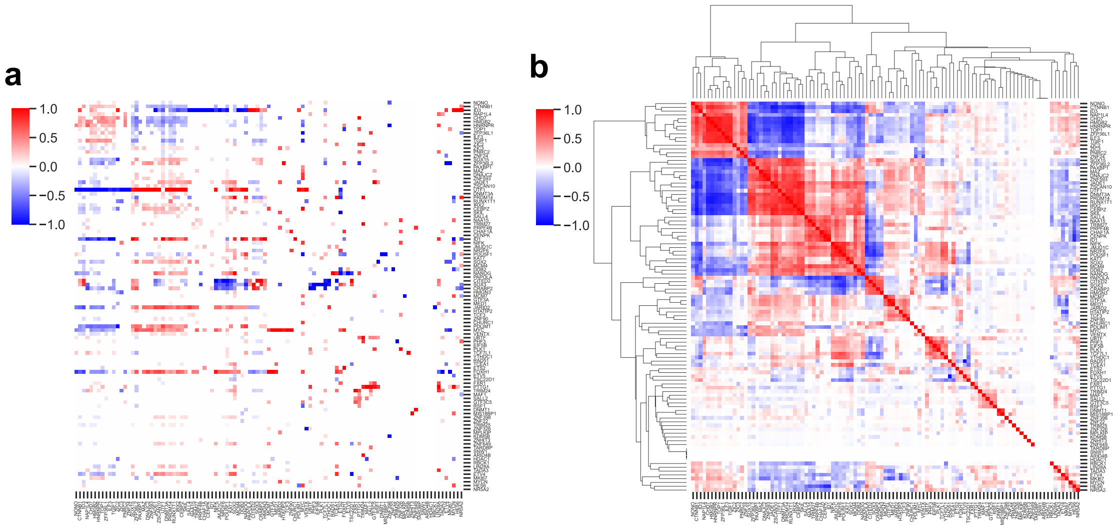


Supplementary Figure 5: **Regulatory coefficients and regulatory correlations in hiPSCs. a**, Regulatory coefficients between 103 genes, showing the strength of regulation from the gene in each column to the gene in each row. The regulatory coefficients with $q$-value $\leq 0.01$ were set to 0, while the others were normalized so that the maximum absolute value of the regulatory coefficients in each column was 1. **b**, Regulatory correlations between 103 genes. A regulatory correlation is the correlation coefficient between the regulatory coefficients of the gene in each column and the gene in each row.

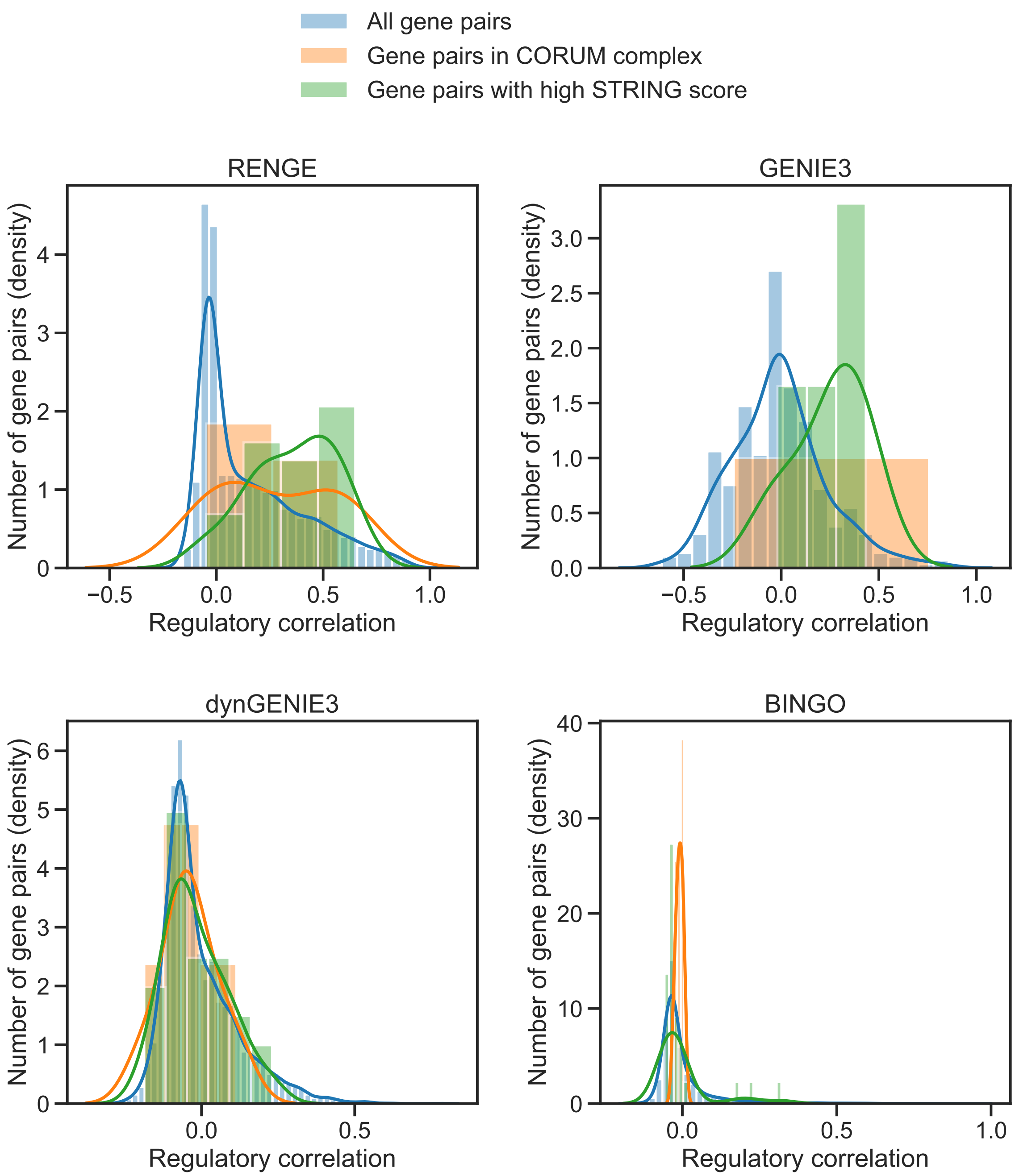


Supplementary Figure 6: **Distributions showing the regulatory correlations among all gene pairs estimated using each method.** Distribution of the regulatory correlations among all possible gene pairs compared with those in the protein complexes obtained from CORUM3.0 and those with high STRING scores. For each method, 1354 regulations, the number of regulations with $q$-value $< 0.01$ in RENGE, were selected in order of confidence level, and the regulatory coefficients for the remaining regulations were set to 0. Regulatory correlations for RENGE were calculated using the absolute value of the regulatory coefficients ignoring the sign for consistency with other methods, which only infer regulatory strength without the sign. Gene pairs with correlation coefficients that could not be calculated due to no inferred regulation from the gene, are not shown. For GENIE3, the kernel density estimate for gene pairs in the CORUM complex is not shown as only one gene pair was included in the CORUM complex.

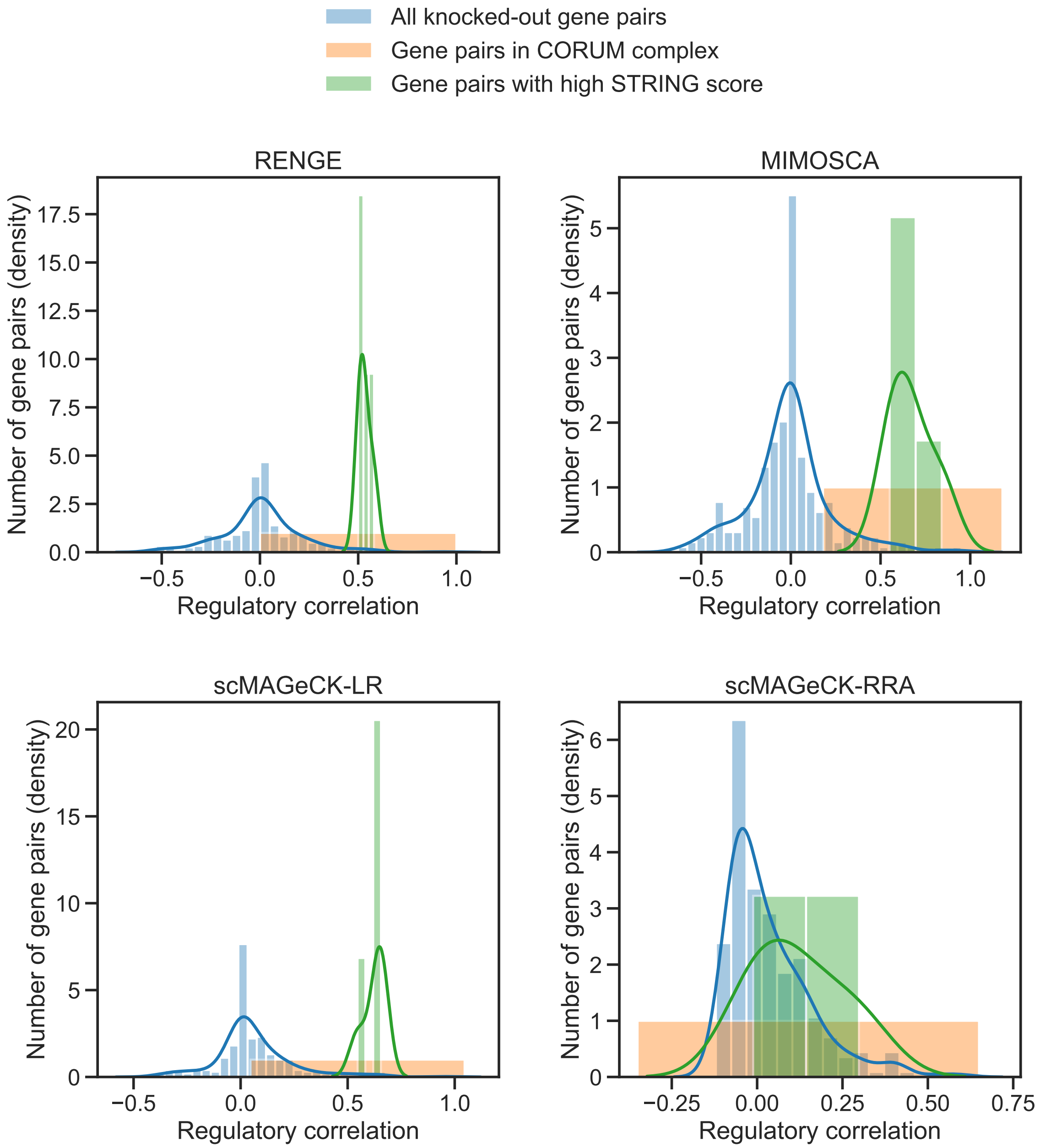


Supplementary Figure 7: **Distributions showing the regulatory correlations among the knocked-out genes that were estimated using each method.** Distribution of the regulatory correlations among all possible pairs in the knocked-out genes when compared with those in the protein complexes obtained from CORUM3.0 and those with high STRING scores. For each method, 346 regulations, the number of regulations with $q$-value $< 0.01$ in RENGE, were selected in order of confidence level of the regulations, and the regulatory coefficients or $q$-values for the remaining regulations were set to 0 or 1, respectively. Gene pairs with correlation coefficients that could not be calculated due to no inferred regulation from the gene, are not shown. For all four methods, kernel density estimates for gene pairs in the CORUM complex are not shown as only one gene pair was included in the CORUM complex.

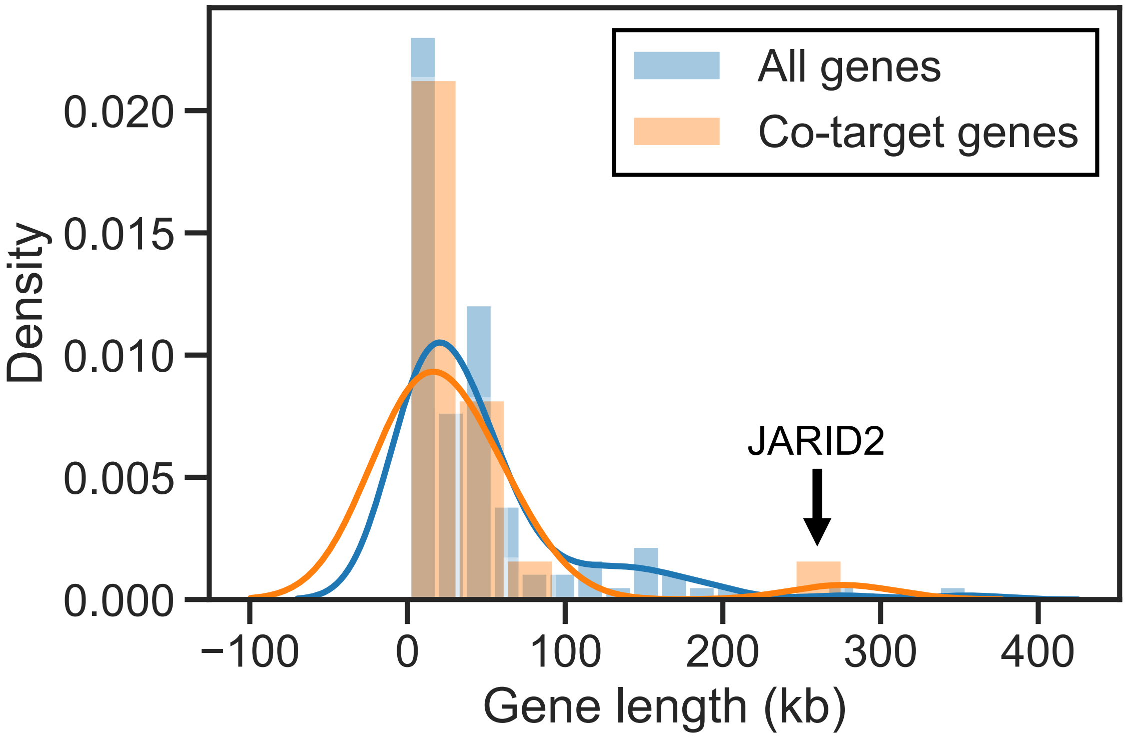


Supplementary Figure 8: **Distribution showing the length of all genes in the GRN versus co-target genes of CHD7 and TOP1.** Co-target genes are those that are regulated by both CHD7 and TOP1 (FDR $< 0.01$). *JARID2*, the longest gene in the co-target genes is identified by an arrow.

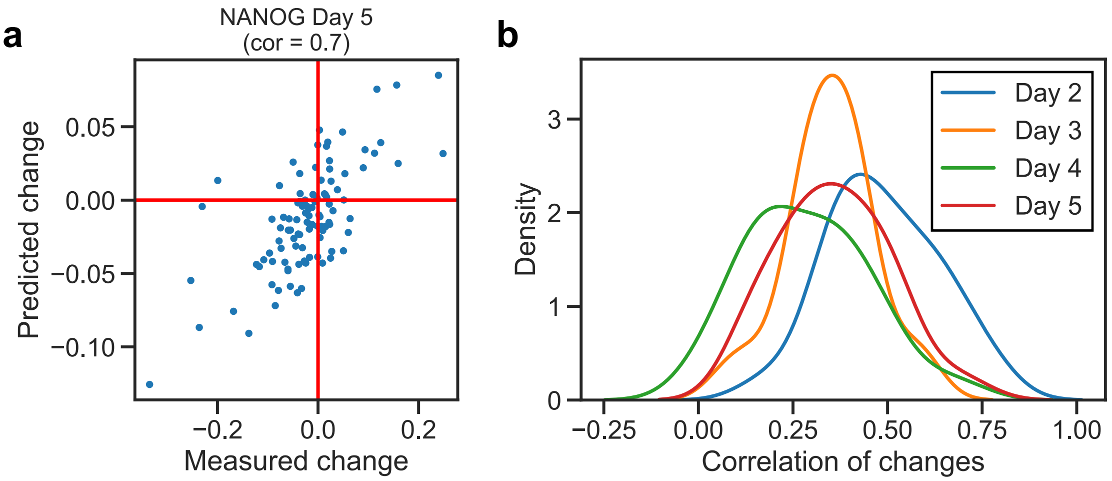


Supplementary Figure 9: **Prediction of expression changes by knockout using RENGE. a**, Relationship between predicted and measured expression changes on day 5 when *NANOG* is knocked out. Each point corresponds to a gene in the network. The horizontal and vertical axes indicate the experimentally measured and predicted changes in expression for each gene when *NANOG* is knocked out, respectively. The prediction was made using the RENGE model learned using data without the *NANOG* knockout. **b**, Distribution of the correlation coefficients between the measured expression changes for each gene and the expression changes predicted by RENGE when each of the 23 genes were knocked out. The RENGE model which was trained without the knockout data for gene $i$ was used to predict the expression changes for each gene when gene $i$ was knocked out.

Supplementary Table 1: gRNA sequences used in the single-cell CRISPR experiment

| Gene symbol | sgRNA ID | Guide sequence |
|---|---|---|
| Safe-cutter control | AAVS1-1 | GTCCCTAGTGGCCCCACTG |
| Safe-cutter control | AAVS1-2 | GGGCCACTAGGGACAGGAT |
| Non-cutter control | CTRL00509 | CGAATTCACTCCCTTAGCG |
| Non-cutter control | CTRL00626 | ACGCCCAGACTGTTAGCGC |
| ETS2 | ETS2_5-2 | TGCAGAGGTTCGGCATGAA |
| ETS2 | ETS2_5-5 | CACAGAATTACCCCAAAGG |
| ETV4 | ETV4_5-1 | GGGCACCCCGGCGCTGGTA |
| ETV4 | ETV4_5-2 | GGTCGGTGCATTTCGAGAG |
| FOXH1 | FOXH1_5-2 | GCTGCACGGCCGGCCATAC |
| FOXH1 | FOXH1_5-4 | CGGTTTCAGATCATCCGTC |
| ID1 | ID1_5-2 | GGCCATCTCGCGCTGCGCC |

| | | |
|---|---|---|
| ID1 | ID1_5-5 | TACATCAGGGACCTTCAGT |
| JARID2 | JARID2_5-4 | ATCGAGTCGGTCCGCGCTC |
| JARID2 | JARID2_5-5 | GGTGATCCCCCCTCCGGAC |
| LIN28A | LIN28A_5-3 | TGTCCATGACCGCCCGCGC |
| LIN28A | LIN28A_5-4 | GAGTAAGCTGCACATGGAA |
| MYC | MYC_2 | TATTTCTACTGCGACGAGG |
| MYC | MYC_7 | GGGTCGATGCACTCTGAGG |
| MYCN | MYCN_5-3 | TCGCGCTTGTTCACGGGAA |
| MYCN | MYCN_5-4 | GGTATTGCCGCCCCAGCCG |
| NANOG | NANOG_g1 | CCTCAGCTACAAACAGGT |
| NANOG | NANOG_g2 | GAGAAGAGTGTCGCAAAAA |
| NR5A2 | NR5A2_5-2 | AGTCCATTGGCTCGGATGA |
| NR5A2 | NR5A2_5-5 | GATGGCCCGGCTAGGAAAG |
| PDLIM1 | PDLIM1_5-1 | GGCAGCAGTCTTTGACTCC |
| PDLIM1 | PDLIM1_Bru | ATGAGCAAGAACTTACCTG |
| POU5F1 | POU5F1_g1 | GTGGGTTTCGGGCACTGC |
| POU5F1 | POU5F1_g15-14 | CTTGCAGGTGGTCCGAGTG |
| PRDM14 | PRDM14_v32_7-4 | GCTCGGTTCCAGTTCACGG |
| PRDM14 | PRDM14_v32_7-6 | GCAGCAGCCACGAGTACGC |
| RUNX1T1 | RUNX1T1 g3 | GAGTTCGCACCCTCGTTCT |
| RUNX1T1 | RUNX1T1 g7-5 | TACCACTAGTCCCAGAACG |
| SOX2 | SOX2_v32_7-4 | GCGGGCGTGAACCAGCGCA |
| SOX2 | SOX2_v32_7-5 | TTATAAATACCGGCCCCGG |
| TRIM24 | TRIM24_4 | TTTGAGCTCACCAGTGGGA |
| TRIM24 | TRIM24_5-4 | TTGCATAAGTTTCATGCGA |
| TRIM25 | TRIM25_5-2 | GGCGCAACAGGTCGCGAAC |
| TRIM25 | TRIM25_v3_6-4 | AACACGGTGCTGTGCAACG |
| VENTX | VENTX_5-4 | TCTATGTCAGGGTTGAGTA |
| VENTX | VENTX_5-5 | GACGTTGAGTAGAAAGCTG |
| ZIC2 | ZIC2_5-4 | CATATTCATGGGGCCGTAC |
| ZIC2 | ZIC2_5-5 | GGGATTGCTCAGTTGCTCG |
| ZIC3 | ZIC3_5-3 | TCGCGCGTTGAGTTGAAGG |

| | | |
|---|---|---|
| ZIC3 | ZIC3_5-5 | ACTGAGCGCCGGCCGCGTA |
| ZNF398 | ZNF398_5-2 | CGTCACTCTCCTTGGACTC |
| ZNF398 | ZNF398_5-5 | ACACCTGCTCGCCAACTGG |
| ZNF649 | ZNF649_5-1 | AATCATGAACAAATGCCTA |
| ZNF649 | ZNF649_5-5 | CACAAACAGGGTATCAAGC |
| ZNF90 | ZNF90_5-3 | TTATGTACATAAAGGATCG |
| ZNF90 | ZNF90_5-5 | TTATGTTTATAAAGGAGTG |

Supplementary Table 2: List of the oligonucleotides used to prepare the gRNA sequencing library

GuideForward

5 ' -AATGATACGGCGACCACCGAGATCTACACTCTTTCCCTACACGACGCTCTTCCGATCT-3 '

GuideReverse-R1

5 ' -CAAGCAGAAGACGGCATACGAGATAGGAGTCCGTCTCGTGGGCTCGGAGATGTGTATAAGAGACAGCGGACTAGCCttatttaaacttgctatgc-3 '

GuideReverse-R2

5 ' -CAAGCAGAAGACGGCATACGAGATCATGCCTAGTCTCGTGGGCTCGGAGATGTGTATAAGAGACAGCGGACTAGCCttatttaaacttgctatgc-3 '

GuideReverse-R3

5 ' -CAAGCAGAAGACGGCATACGAGATGTAGAGAGGTCTCGTGGGCTCGGAGATGTGTATAAGAGACAGCGGACTAGCCttatttaaacttgctatgc-3 '

GuideReverse-R4

5 ' -CAAGCAGAAGACGGCATACGAGATCAGCCTCGGTCTCGTGGGCTCGGAGATGTGTATAAGAGACAGCGGACTAGCCttatttaaacttgctatgc-3 '

Supplementary Table 3: Genes with ChIP-seq data obtained from the ChIP-Atlas.

| Gene with ChIP-seq | Knock out |
|---|---|
| FOXH1 | ✔ |
| JARID2 | ✔ |
| MYC | ✔ |
| NANOG | ✔ |
| NR5A2 | ✔ |
| POU5F1 | ✔ |
| PRDM14 | ✔ |
| RUNX1T1 | ✔ |
| SOX2 | ✔ |
| ZNF398 | ✔ |
| CHD7 | |
| CTNNB1 | |
| DNMT1 | |
| DNMT3A | |
| KDM5B | |
| MED1 | |
| SALL4 | |
| TCF3 | |
| TCF7L1 | |
| UBTF | |

Supplementary Table 4: Correlation coefficients between the confidence of inferred regulation and ChIP-seq $-10 * \log_{10}(\text{MACS2 } q\text{-value})$

| | Regulations by KO genes | Regulations by non-KO genes |
|---|---|---|
| RENGE | 0.268 | **0.313** |
| GENIE3 | **0.270** | 0.078 |
| dynGENIE3 | 0.106 | 0.091 |
| BINGO | 0.034 | 0.049 |
| MIMOSCA | 0.205 | - |
| scMAGeCK-LR | 0.230 | - |
| scMAGeCK-RRA | 0.080 | - |

Supplementary Table 5: Correlation coefficients between the measured and predicted expression changes by RENGE for each day when each of the 23 genes were knocked out.

| KO gene | Day 2 | Day 3 | Day 4 | Day 5 |
|---|---|---|---|---|
| ETS2 | 0.78 | 0.42 | 0.44 | 0.39 |
| ETV4 | 0.43 | 0.08 | 0.18 | 0.13 |
| FOXH1 | 0.46 | 0.37 | 0.21 | 0.20 |
| ID1 | 0.51 | 0.42 | 0.30 | 0.38 |
| JARID2 | 0.51 | 0.30 | 0.14 | 0.30 |
| LIN28A | 0.51 | 0.34 | 0.18 | 0.35 |
| MYC | 0.62 | 0.27 | 0.17 | 0.15 |
| MYCN | 0.73 | 0.59 | 0.34 | 0.25 |
| NANOG | 0.56 | 0.34 | 0.67 | 0.70 |
| NR5A2 | 0.62 | 0.56 | 0.34 | 0.52 |
| PDLIM1 | 0.18 | 0.14 | 0.06 | 0.29 |
| POU5F1 | 0.67 | 0.29 | 0.50 | 0.46 |
| PRDM14 | 0.33 | 0.45 | 0.50 | 0.56 |
| RUNX1T1 | 0.34 | 0.27 | 0.22 | 0.39 |
| SOX2 | 0.41 | 0.35 | 0.44 | 0.52 |
| TRIM24 | 0.59 | 0.41 | 0.21 | 0.50 |
| TRIM25 | 0.34 | 0.35 | 0.34 | 0.42 |
| VENTX | 0.42 | 0.29 | 0.36 | 0.26 |
| ZIC2 | 0.46 | 0.25 | 0.09 | 0.14 |
| ZIC3 | 0.40 | 0.31 | 0.01 | 0.24 |
| ZNF398 | 0.37 | 0.41 | 0.15 | 0.36 |
| ZNF649 | 0.35 | 0.42 | 0.32 | 0.46 |
| ZNF90 | 0.65 | 0.42 | 0.41 | 0.31 |

Supplementary Table 6: dyngen parameters used for the benchmark.

| Parameter | Value |
|---|---|
| total_time | simtime_from_backbone(backbone) + 100 |
| num_timepoints | ceiling(total_time / 100 × 4) |
| timepoint (of KO) | (total_time - 100) / total_time |
| multiplier | rbinom(num_simulations, 1, 0.3) |
| pct_between | 0.1 |
| num_cells | 25 × num_timepoints |
| num_tfs | 100 |
| num_targets | 0 |
| num_hks | 0 |
| n_census | 10 × num_cells |
| census_interval | floor((total_time × num_simulations) / n_census) |
| tau | 0.5 |
